# MitoCore: A curated constraint-based model for simulating human central metabolism

**DOI:** 10.1101/141101

**Authors:** Anthony C. Smith, Filmon Eyassu, Jean-Pierre Mazat, Alan J. Robinson

## Abstract

**Background:** The complexity of metabolic networks can make the origin and impact of profound changes in central metabolism occurring during disease difficult to understand. Computer simulations can help unravel this complexity, and progress has advanced in genome-scale metabolic models. However, many current models produce unrealistic results when challenged to simulate abnormal metabolism as they include incorrect specification and localization of reactions and transport steps, incorrect reaction parameters, and confounding of prosthetic groups and free metabolites in reactions. Other common drawbacks are due to their scale, such as being difficult to parameterise and simulation results being hard to interpret. Therefore, it remains important to develop smaller, manually curated models to represent central metabolism accurately.

**Results:** We present MitoCore, a manually curated constraint-based computer model of human metabolism that incorporates the complexity of central metabolism and simulates this metabolism successfully under normal and abnormal conditions, including hypoxia and mitochondrial diseases. MitoCore describes 324 metabolic reactions, 83 transport steps between mitochondrion and cytosol, and 74 metabolite inputs and outputs through the plasma membrane, to produce a model of manageable scale for easy data interpretation. Its key innovations include accurate partitioning of metabolism between cytosol and mitochondrial matrix; correct modelling of connecting transport steps; proper differentiation of prosthetic groups and free co-factors in reactions; and a new representation of the respiratory chain and the proton motive force. MitoCore’s default parameters simulate normal cardiomyocyte metabolism, and to improve usability and allow comparison with other models and types of analysis, its reactions and metabolites have extensive annotation, and cross-reference identifiers from Virtual Metabolic Human database and KEGG. These innovations—including over 100 reactions absent or modified from Recon 2—are essential to model central metabolism accurately.

**Conclusion:** We anticipate MitoCore as a research tool for scientists, from experimentalists looking to interpret data and generate further hypotheses, to experienced modellers predicting the consequences of disease or using computationally intensive methods that are infeasible with larger models, as well as a teaching tool for those new to modelling and needing a small manageable model on which to learn and experiment.

## Background

Human central metabolism is a large and complex system under sensitive homeostatic control, and its disturbance is causative or associated with many diseases and responses to toxins. However, it is often difficult to relate more than a handful of these changes to their underlying origin or their down-stream impact, due to the highly connected nature of the reactions of central metabolism. Computer models are widely accepted in many fields as a tool to incorporate complexity and simulate changes, allowing predictions to be made and providing a unifying framework to interpret empirical data, especially from large, noisy and incomplete data sets. Yet modelling is treated with scepticism by many biomedical researchers despite their potential broad utility [1]. Simple models of enzyme kinetics (using the assumptions of Henri-Michaelis-Menten kinetics [2]) are familiar to biomedical scientists, but are impractical for simulations of central metabolism due to every reaction needing parameterisation, alongside the computational expense of solving the large set of differential equations. However, constraint-based models of metabolism used in conjunction with methods such as flux balance analysis [3] are particularly useful for simulating metabolic changes in large metabolic networks as they can incorporate flexibility, do not require any kinetic parameters and are computationally inexpensive. Many genome-scale constraint-based models [4–10], representing known enzymes encoded in the human genome, have covered central metabolism and used to successfully model some diseases [11,12]. But these models do not simulate the realistic production rate of ATP (with one recent exception [10]), a crucial element of modelling central metabolism. Furthermore, the interpretation of simulation results from thousands of reactions is difficult (especially for new-comers), and erroneous “short-circuits” and energy-generating cycles commonly occur without extensive manual curation [13]. In addition, attempts to simulate disease can result in the prediction of physiologically improbable reaction fluxes. These are caused by several common problems including: incorrect parameters for directionality constraints, the assignment of reactions to the wrong cellular compartments, or inaccurate representations of pathways, enzymes, transport steps, prosthetic groups and metabolites. These errors can introduce unrealistic bypasses and shuttles that appear to compensate for a disease state. For example, proton-coupled mitochondrial transporters running in reverse and thus pumping protons that contribute to ATP generation, and the confounding of free co-factors with prosthetic groups, especially the FAD of mitochondrial succinate dehydrogenase and electron-transferring flavoprotein (ETF), leading to unrealistic fluxes of electrons between isolated complexes. These problems are common in genome-scale models that include an initial auto-generation of the reaction network from databases that can include incomplete or incorrect annotation. These issues are particularly acute for modelling mitochondrial metabolism and metabolite transport, as all the current genome-scale models neglect the electrical gradient component (ΔΨ) of the proton motive force (PMF) and the correct proton cost of making ATP by ATP synthase in animals [14]. It is also sometimes questionable whether the enormous size and complexity of these genome-scale models benefits simulations of subsystems of cellular metabolism, such as central metabolism. Furthermore, genome-scale models make some techniques computationally infeasible due to their scale, such as elementary mode analysis [15].

By using smaller curated models validated against data from normal and disease metabolism these problems can be avoided. A more focussed and carefully defined model also allows the user to be confident in each reaction and more clearly elucidate the behaviour of the system, including any short-comings, and interpret the results more readily. We previously applied this approach for our *i*AS253 model of the mitochondrion, which we used to simulate metabolic diseases of the tricarboxylic acid cycle [16]. This model was then used as a basis to simulate other disorders including hypoxia during cardiac ischemia [17], fumarate hydratase deficiency [18] and common diseases of the mitochondrial electron transport chain [19]. These simulation results were used to generate detailed mechanistic hypotheses for data interpretation and to design further experiments. However, we recognised that this model could be improved upon by constructing a new model that encompassed more of central metabolism and explicitly modelled physiochemical features such as the mitochondrial proton motive force. In particular, ease of use would be improved by providing extensive annotation of reactions and their parameters.

Here we present MitoCore, a new constraint-based model of central metabolism that addresses these issues and comprehensively expands upon and refines our previous mitochondrial models. This model has been designed to be easy-to-use, includes extensive annotation, has default parameters to simulate human cardiomyocyte metabolism, and is encoded in the widely used SBML format [20]. We anticipate the model will be of great use to those wishing to interpret empirical data by comparing it to simulations of central metabolism and thus investigate predictive models of disease and toxicology.

## Results

### Building the MitoCore model of human central metabolism

MitoCore is a constraint-based model of central metabolism with two compartments; one representing the cytosol, outer mitochondrial membrane, inter-membrane space and cytosolic side of the inner mitochondrial membrane, and the other the mitochondrial side of the inner membrane and the mitochondrial matrix. It models 324 reactions of which 157 have been assigned to the mitochondrial compartment, and 167 assigned to the cytosolic compartment. 83 mitochondrial transport steps connect the two compartments, of which 30 are modelled on the known transport mechanisms of characterised transport proteins of the inner mitochondrial membrane, whereas the other 53 represent known transport capabilities of the membrane, such as diffusion of small metabolites. 74 transport steps at the cytosolic boundary represent the import and export of metabolites across the plasma membrane, such as oxygen, carbon dioxide, glucose, fatty acids and amino acids. For convenience MitoCore includes four ‘pseudo’ reactions that summarise aspects of its biological activity and can be used by flux balance analysis as objective functions: ATP hydrolysis (representing cellular ATP demand), and the biosynthesis of heme, lipids and amino acids (representing the anabolism of biomolecules for cellular growth).

To create the MitoCore model of central metabolism, we built a list of candidate reactions to include by considering human reactions in the KEGG [21], HumanCyc [22] and BRENDA [23] databases that use any metabolites involved in central metabolism, and assigned each reaction to the appropriate cellular compartment(s) by assessing the localisation evidence collated in the MitoMiner database [24] for its catalysing protein. Reaction directionality was assigned by taking the consensus from annotation in metabolic databases, estimates of Gibbs free energy [25,26], and general rules of irreversibility [27]. For each reaction extensive additional annotation was recorded including the original KEGG identifiers, EC number, description, gene mappings (both HUGO gene symbol and Ensembl identifiers), and evidence for the gene’s expression in heart and the protein’s mitochondrial localisation.

To enable comparison of results from MitoCore to those of the popular genome-scale models such as Recon 2 [5], MitoCore re-used identifiers for metabolites and reactions present in the Virtual Metabolic Human database (https://vmh.uni.lu) where possible. However, it was necessary to create 105 new reactions for MitoCore (additional file 6) that were either absent from the Virtual Metabolic Human database (such as transport steps or compartment specific versions), were inaccurately described (such as specifying prosthetic FAD as a free cofactor), or were needed to represent new features (such as the proton motive force). New reaction identifiers were appended with the suffix ‘MitoCore’. Pathways represented in MitoCore include glycolysis, pentose phosphate pathway, TCA cycle, electron transport chain, synthesis and oxidation of fatty acids, ketone body and amino acid degradation and cover all parts of central metabolism involved directly or indirectly with ATP production.

Finally, MitoCore was extensively tested to ensure that it contained no erroneous energy-generating cycles, it was capable of simulating disorders such as ischemia and mitochondrial diseases, and that each reaction was capable of having a flux (depending on the constraints placed on the cytosolic boundary transport steps) and so ensure it contained no reaction dead-ends. MitoCore was encoded in SBML [20] (additional file 1) and a companion annotation Excel spreadsheet produced (additional file 2).

#### Simulating cardiomyocyte metabolism using MitoCore and flux balance analysis

MitoCore’s default reactions and parameters are optimised for cardiomyocytes and use the metabolites available to healthy hearts of glucose, fatty acids, ketone bodies and amino acids (references listed in additional file 2). To demonstrate that MitoCore was capable of producing physiological relevant results using these parameters, we simulated cardiomyocyte metabolism by using flux balance analysis [3]. Flux balance analysis calculates the optimum rate of turnover, or flux, of metabolite through each reaction of a network, given a particular objective. To reflect the primary role of central metabolism in cardiomyocytes we set the simulation’s objective as maximum ATP production (by maximising flux through the pseudo reaction of ATP hydrolysis) and calculated the optimum reaction fluxes through central metabolism. The resultant reaction fluxes simulated core metabolism correctly with activity of all respiratory complexes and the TCA cycle (Fig 1, additional file 2). As metabolic fuels were provided in slight excess, the availability of oxygen limited the overall fluxes. Simulated ATP production was 100.9 μmol/minute/gram of dry weight. Sources of acetyl-CoA for the TCA cycle were fatty acid degradation (55.0%), glucose oxidation (26.4%), lactate oxidation (8.4%), ketone body degradation (6.1%), amino acid degradation (3.8%) and glycerol oxidation (0.3%). The amino acids degraded and used to produce ATP were histidine, isoleucine, leucine, lysine, threonine, valine, arginine, aspartate, cysteine, glycine, proline, serine, asparagine, and alanine. Ammonia, produced as a by-product of amino acid degradation, was exported from the system.

**Fig 1.**
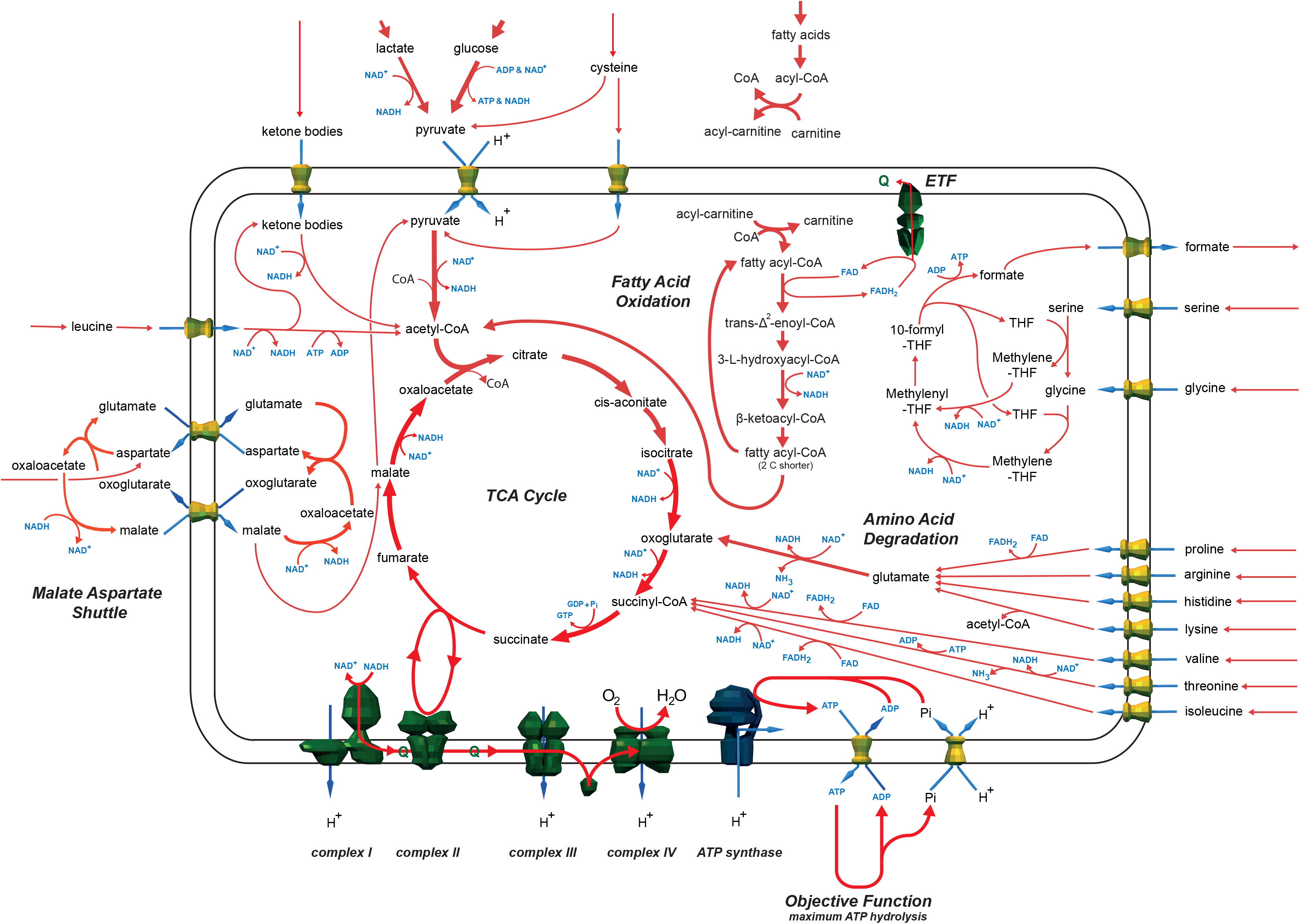
Summary of the major active pathways of central metabolism in the flux balance analysis simulation using the MitoCore model with its default parameters and the objective function of maximum ATP production. Values of all fluxes are reported in additional file 1.

### Metabolite degradation and ATP yields using MitoCore

To demonstrate how different metabolites are degraded appropriately in the MitoCore model and the effect of the implementation of the proton motive force on ATP production, we performed a series of simulations using particular ‘fuel’ metabolites in isolation. We simulated the oxidation and degradation of glucose, lactate, hexadecanoic acid, hydroxybutanoate, acetoacetate and 20 different amino acids in separate simulations. In each simulation, one fuel was allowed a maximum uptake of 1 μmol/minute/gram of dry weight while all others were set to zero, then the objective function was set to maximise ATP hydrolysis, and the optimum reaction fluxes in the metabolic network were calculated by using flux balance analysis (additional file 3). The simulations showed unsurprisingly that fatty acids were the most energy rich fuels to metabolise (ATP production of 112 μmol/minute/gram of dry weight), followed by the amino acids tryptophan (43 μmol/minute/gram of dry weight), isoleucine (38 μmol/minute/gram of dry weight), leucine (37 μmol/minute/gram of dry weight) and phenylalanine (36 μmol/minute/gram of dry weight) and then glucose (33 μmol/minute/gram of dry weight).

### The effect of proton leak on the electron transport chain in MitoCore

To show the MitoCore model can simulate experimentally induced conditions that are directly affected by the proton motive force, we performed a series of simulations representing an increasing proton leak across the mitochondrial inner membrane. We introduced a flux through the transport step that represents the leak of matrix protons to the cytosol through the uncoupling protein 2 (UCP2, Reaction ID: HtmB_MitoCore) in cardiomyocytes [28], while using the objective function of maximum ATP hydrolysis during flux balance analysis. The optimum reaction fluxes (additional file 4) were identical to those under default constraints with the exception of the steps involved in mitochondrial ATP synthesis. The flux through ATP synthase was progressively reduced as the proton leak increased until it reversed under very high leak, using ATP synthesized in the Krebs cycle. At the highest level of proton leak the ATP/ADP carrier reversed to provide additional ATP from glycolysis (Reaction ID: ATPtmB_MitoCore). Similarly the phosphate carrier also reversed at the highest proton leak value.

### Discussion

Here we present MitoCore, a curated, constraint-based model of human central metabolism (additional file 1) designed as a predictive model of metabolism in disease and toxicology, and for use by a wide range of researchers. It covers all major pathways involved in central metabolism using 407 reactions and mitochondrial transport steps, and 74 transport steps over the plasma membrane. To increase the metabolic flexibility of MitoCore, we included a large number of reactions that were not assigned to classical pathways, but could have potentially important roles in supporting central metabolism. MitoCore was parameterised and annotated for cardiomyocyte metabolism, which is useful for many types of analyses as the cardiomyocyte can metabolise a wide range of substrates and has reactions common to many other cell types, as well as representing the metabolism of an organ of utmost importance in human health, disease and toxicology. This allows the simulation results to be generalisable, without having features that are particularly cell specific, such as those found only in hepatocytes. However, we have included reactions that are switched off by default in the cardiac model, but can be activated to represent the abilities of other cell types, e.g. gluconeogenesis, ketogenesis, β-alanine synthesis and folate degradation. Thus, the model allows biologically relevant flux distributions to be generated ‘out of the box’ without altering the model, while allowing for easy modification to represent other cell types.

MitoCore has several unique features. The first is the accurate partitioning of metabolism between the mitochondrion and the cytosol by using extensive localisation data and annotation. This partitioning has an important impact on model behaviour as the limited transport steps into the mitochondrial matrix result in dramatic differences in metabolite availability in the matrix compared to the cytosol. Therefore, it is crucial that reactions are assigned to the correct compartment. We achieved this by manually evaluating the localisation of each reaction’s catalysing protein by using the mitochondrial localisation evidence in the MitoMiner database [24]. MitoMiner collates GFP tagging, large-scale mass-spectrometry mitochondrial proteomics studies and mitochondrial targeting sequence predictions, with detailed annotation from the Gene Ontology and metabolic pathway data from KEGG. MitoMiner also contains homology information allowing localisation evidence to be shared amongst species. For reactions catalysed by enzymes with a large amount of evidence for mitochondrial localisation but lacking specific evidence for being in the mitochondrial matrix or matrix side of the inner membrane, we applied the principle of metabolite availability—a reaction can only be present in a compartment if all its substrates are available and its products can be used by reactions within the same compartment [16]. A summary of this localisation evidence is provided in the mitochondrial evidence section of the supplementary annotation file (additional file 2, ‘Reaction & Fluxes’ worksheet) and consists of confidence scores from the MitoCarta 2 [29] inventory of genes that encode mitochondrial proteins. This dataset is derived by using machine learning to evaluate mitochondrial localisation data from training on collections of characterised mitochondrial and non-mitochondrial genes.

The partitioning of metabolism between the mitochondrion and the cytosol logically led us to reconsider the proton motive force (PMF) and the role of protons crossing the inner mitochondrial membrane as part of oxidative phosphorylation, as it produces the majority of cellular ATP. The PMF is achieved by complexes I, III and IV pumping matrix protons across the mitochondrial inner membrane and into the intermembrane space. This proton-pumping creates a proton motive force (PMF) across the membrane that has two components: a proton gradient (ΔpH) coupled with an electrical membrane potential (ΔΨ). The energy for proton-pumping comes from the transfer of electrons down the respiratory chain from NADH and ubiquinone to oxygen, to form water. Additional electrons are passed into the respiratory chain from the TCA cycle by complex II, and from the degradation of fatty acids and amino acids by the electron-transfer flavoprotein (ETF). ATP synthase uses the PMF to power ATP synthesis from ADP and phosphate by channelling protons back across the membrane. It is thus clear that it is necessary to distinguish as Peter Mitchell did [30], the protons involved in chemical reactions taking place in an isolated compartment (“scalar protons”), from the protons crossing between compartments (“vectorial protons”). To better model oxidative phosphorylation, we devised a new representation of the PMF and mitochondrial respiratory chain—the second unique feature of MitoCore. MitoCore’s representation of the respiratory chain differs in many key aspects to other metabolic models, due to how it models vectorial protons and accounts for both components of the PMF. MitoCore represents the PMF as a metabolite that is co-transported in steps that transport charged metabolites or protons across the inner mitochondrial membrane, such as the reactions of the respiratory complexes, and in proton-coupled and electrogenic transport steps. As MitoCore’s representation of the PMF incorporates both ΔΨ and ΔpH, we reflect their relative contributions to the overall PMF by co-transporting 0.82 PMF metabolites for transport steps that affect ΔΨ and 0.18 PMF metabolites for transport steps that affect ΔpH. These values are the average of published figures of the relative contributions of ΔpH and ΔΨ by several authors (see additional file 5 for details and references). Therefore, the reactions that represent complexes I, III and IV of the respiratory chain move PMF metabolites that correspond to the number of protons they pump from the matrix to the cytosol, as a proton in this case affects both ΔΨ and ΔpH. ATP synthase needs to transport 2.7 PMF metabolites back to the matrix to synthesise one molecule of ATP (as it uses 2.7 protons per molecule of ATP [14]). Transport steps between the two compartments are modelled in the same way; for example the mitochondrial ATP/ADP carrier 1 (SLC25A4) requires 0.82 PMF to be co-transported with each imported ADP^3-^ nucleotide to reflect the charge difference between ATP^4-^ and ADP^3-^ that affects only ΔΨ (equation 1), whereas the proton-coupled phosphate carrier (SLC25A3) imports 0.18 PMF as overall transport is electro-neutral and so only affects ΔpH (equation 2):

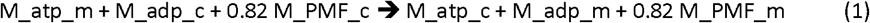

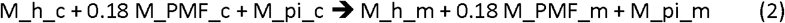

This separate modelling of vectorial protons using an additional new PMF metabolite enabled the impact of electrogenic transporters on the PMF to be accounted for in flux balance analysis simulations for the first time and prevented simulation artefacts where (scalar) protons generated or removed from other parts of metabolism allow unrealistically high ATP production, as can be the case in other models. We have removed FADH from reactions (as specified in databases such as KEGG) where it is treated as a redox cofactor (such as for complex II and the ETF), and instead directly coupled the reactions to the reduction of ubiquinone to ubiquinol. This keeps separate the electrons entering the respiratory chain from different sources, which can otherwise become connected via a shared FADH metabolite, and is particularly relevant under perturbed conditions where the erroneous connection of free co-factor and bound prosthetic FADH can cause unrealistic bypasses to occur. This problem is endemic in large-scale models that auto-generate reaction networks directly from metabolic databases without manual curation of reactions. We model the generation of reactive oxygen species (ROS) at 0.001% of the flux through complex I, to reflect the primary site of ROS generation from the respiratory chain and reduce the efficiency of proton pumping as electrons escape [31,32]. Further reactions convert ROS to water at the expense of NADPH.

The third unique feature of MitoCore is the modelling of the transport steps that connect the cytosolic and mitochondrial compartments. We included four different transport categories. First were transport steps based on the characterised mitochondrial transport proteins. Many carriers can transport a range of related substrates (although with different affinities) and each metabolite combination was modelled as a separate step including counter exchange and proton-coupling. In some cases proton-coupled transporters were represented by two reactions to model separately the forward and reverse directions, and a proton (plus cotransported PMF metabolite) only used for movement down the proton gradient. This prevented transporters being used in reverse to pump protons artificially (and so transferring PMF) and thereby contribute to ATP production by the ATP synthase. The second category was for metabolites whose transporters are unidentified. We chose to model these steps as uniporters that were not proton coupled. If the metabolite was charged and moving into the matrix we assumed this would impact ΔΨ and co-transported 0.82 PMF metabolite accordingly. The third category was for metabolites that can diffuse across the inner mitochondrial membrane—including oxygen, carbon dioxide and water—and these were modelled as reversible uniport transport steps. Finally we modelled the insertion of lipids into the mitochondrial inner membrane via the flippases, which was coupled to ATP hydrolysis.

Other improvements to MitoCore included modelling the protein complexes as one reaction rather than a series of linked reactions, as can often be found in metabolic pathway databases and other models. This is particularly important for gene knockout simulations where a gene deficiency will knock out a protein complex rather than just one of the constituent reactions. To facilitate gene-based analyses (such as knock out studies) we provide reaction-to-gene mappings with both gene symbols and Ensembl identifiers. To summarise the behaviour of MitoCore, we also defined four ‘pseudo’ reactions for biosynthesis of biomolecules (amino acids, lipids, DNA) required by cells for growth and ATP hydrolysis to reflect the energy demand of the cell. These pseudo reactions are designed to be used as objective functions during flux balance analysis.

A weakness of many metabolic models is the lack of provenance about their components. For example recording why reactions have been included, why directionality constraints have been set, or the origin of reaction parameters. Thus for MitoCore we created a supplementary annotation spreadsheet (additional file 2) to record this provenance. For example, the directionality evidence section describes why a constraint has been set; a manually evaluated consensus of information from the KEGG [21], HumanCyc [22] and BRENDA [23] databases, general rules of irreversibility [27], large ΔG values from eQuilibrator [26] or estimated using a group contribution method [25], and information from the literature. In cases for which reaction directionality was unclear, it was kept reversible. To address why a reaction has been included in our cardiomyocyte model, we included a heart expression section consisting of RNAseq and immunochemistry expression levels of genes taken from the Human Protein Atlas (version 14) [9]. The spreadsheet also includes gene mappings, identifiers from Recon 2 and KEGG, mitochondrial localisation evidence (as described above) and baseline reaction fluxes when the objective function was maximum ATP production under normal conditions. The spreadsheet also serves as a useful template to map flux distributions onto, as reaction fluxes can be grouped in an intuitive way directly against useful supplementary information. When combined with the small size of the model, simulations can be generated extremely quickly and then easily interpreted.

To demonstrate that MitoCore functions realistically, we simulated cardiomyocyte metabolism using the default parameters with flux balance analysis [3] and the objective of maximum ATP production. The reaction fluxes showed central metabolism was modelled correctly, with the largest fluxes through the TCA cycle and respiratory chain. Numerous fuel sources (fatty acids, glucose, lactate, ketone bodies and amino acids) were imported, degraded and entered the TCA cycle at several different points (Figure 1). The sources of acetyl-CoA for the TCA cycle were similar to the experimental measurements of well-perfused heart [33]— experiments report 60–90% of acetyl-CoA derives from fatty acids compared to 55% in the MitoCore simulation, whereas glycolysis (including lactate oxidation) accounts for 10–40% of acetyl-CoA compared to 34.8% in the MitoCore simulation. These results support the biological relevance and parameter choice of the model.

To show the importance of explicitly modelling the PMF, we simulated the maximum ATP production achievable using 1 μmol/minute/gram of dry weight of common metabolic fuels in isolation (effectively calculating their ATP yields) including for glucose and the fatty acid hexadecanoic acid (additional file 3). In MitoCore each glucose produced 33 ATP in comparison to 32 calculated theoretically [34] and 32 in Recon 2.2 [10]. For the fatty acid hexadecanonic acid 112 ATP was produced compared to 108 theoretically [34] and 107 in Recon 2.2. A significant difference between the models is the number of protons required by ATP synthase to produce one molecule of ATP—MitoCore uses 2.7 (based on the structure of the ATP synthase [14]) whereas the theoretical calculations use 2.5 and Recon 2.2 uses 4.0. Due to also considering additional factors that affect ATP production—including the impact of all the transport steps on the PMF as well as ROS production and removal—we believe our figure is likely to be more accurate than these other calculations. When comparing maximum ATP production using default parameters to the previous iAS253 model [16], ATP production was notably lower (101 vs 140 μmol/minute/gram of dry weight) demonstrating that the separation between the PMF and protons is an important improvement. These simulations also show that the model in each case uses canonical degradation pathways, further increasing confidence that the model is capable of producing realistic results using a wide range of metabolites. The results from the proton leak simulations (additional file 4) show the use of metabolites to represent the PMF enables the model to replicate experimental observations under perturbed conditions that would otherwise be impossible, such as the reversal of ATP synthase, the ATP/ADP carrier and the phosphate carrier, which is well-known behaviour in absence of respiratory chain activity, for instance in ρ° cells to maintain a proton motive force [35,36].

## Conclusions

MitoCore is a constraint-based model of human central metabolism, provided with default parameters that provide physiologically realistic reaction fluxes for cardiomyocytes. The model has several innovations including a new representation of the respiratory chain and proton motive force, and partitioning of reactions to subcellular compartments based on the latest localisation evidence. Each of the 402 reactions was manually evaluated for directionality, expression of its gene in heart and subcellular localisation of its protein resulting in an accurate depiction of central metabolism. To allow MitoCore to be easily used and to make the results directly comparable with other models and compatible with other types of analyses, we have used identifiers from the Virtual Metabolic Human database for both reactions and metabolites where possible, and recorded KEGG identifiers in the annotation. To help ease of use, we also provide an annotation spreadsheet that provides gene mappings, localisation and heart expression evidence, and notes on parameter choice. MitoCore is provided in SBML format to be compatible with a wide range of software. We hope MitoCore will be of use as a research tool to a wide range of biomedical scientists and students—from experienced modellers interested in central metabolism or using computationally intensive methods that are infeasible on a genome-scale model, to those new to modelling who would like begin by using a small manageable model, with application as a predictive model of disease.

## Methods

### Identifying reactions of central metabolism to include in MitoCore

An updated version of the iAS253 mitochondrial model was the starting point for the MitoCore model [19]. The model was expanded by searching for human reactions in KEGG [21], HumanCyc [22] and BRENDA [23] that were missing from the iAS253 model and could impact central metabolism. Each reaction was reassessed for subcellular localisation and directionality (see below), and to ensure the reactions were present in most tissues including cardiac, tissue expression of the genes and proteins was verified by using the Human Protein Atlas [9].

### Partitioning reactions between the cytosol and mitochondrion

To partition the reactions into either cytosol or mitochondrion, each enzyme was manually evaluated using the mitochondrial localisation evidence in the MitoMiner database [24]. All available experimental localisation evidence, mitochondrial targeting sequence predictions and annotation were considered, including from homologs from mouse, rat and yeasts. For reactions with strong evidence for mitochondrial localisation, but where matrix localisation is unclear, we used the principal of metabolite availability [16]. Reactions residing in the mitochondrial matrix or matrix side of the inner membrane were assigned to the mitochondrial compartment. Transport steps were created to connect the cytosol and matrix based on the transport properties of the membrane (active transport, diffusion, etc.). Each reaction was cross-referenced with the Virtual Metabolic Human database (https://vmh.uni.lu) and used its identifiers for reactions and metabolites where possible.

### Assigning reaction directionality

Reaction directionality was manually evaluated for each reaction in MitoCore. The KEGG [21], HumanCyc [22] and BRENDA [23] databases were consulted and general rules of irreversibility were taken into account (such as most reactions consume ATP rather than produce it and carbon dioxide is normally produced not consumed) [27]. We also considered the ΔG values for reactions, both calculated by eQuilibrator [26] and estimated using the group contribution method [25], and large changes noted. Finally we consulted the literature if reaction directionality was conflicted or unclear. If support for irreversibility was poor or a consensus could not be found, the reaction was assigned reversible. The information used to make each assessment was recorded in the directionality evidence column of the annotation spreadsheet (additional file 2). Further refinement of reaction directionality was used to eliminate loops that could produce metabolites such as ATP and NADH for ‘free’, and to prevent the interconversion of NADH and NADPH unless experimentally verified.

### Defining reactions

To improve modelling of the proton motive force (PMF) across the mitochondrial inner membrane, we introduced a pseudo-metabolite to model the effect on the PMF of proton and electrogenic transport steps across the inner mitochondrial membrane. Reactions representing respiratory complexes as well as proton-coupled transport steps were rewritten to use this new species. To prevent unrealistic bypasses between free and complex-bound prosthetic flavin adenine dinucleotides FADH and FAD+ were removed from all reactions and replaced with ubiquinone and ubiquinol. Many reactions were rewritten to represent better the underlying biology, such as the generation of ROS from complex I and the proton-pumping stoichiometry of the respiratory complexes. In some cases the subcellular localisation was changed by using mitochondrial metabolite species, or new reactions were written using existing metabolite species.

In total 105 MitoCore reactions were different to or absent from Recon 2 (additional file 6). To highlight where there are differences between MitoCore reactions and corresponding reactions in the Virtual Metabolic Human database, the MitoCore reaction identifiers use the suffix ‘MitoCore’ and the same Recon 2 identifier if one exists.

### Refining the MitoCore model

Finally, the model was extensively tested to ensure all reactions were capable of having flux, erroneous energy-generating cycles were removed, the model behaved physiologically under normal and perturbed conditions, and all the objective functions were feasible. The model was encoded in SBML v2.1 and its validity checked with the SBML online validator [37].

### Simulating cardiomyocyte metabolism

We simulated metabolism in cardiomyocytes with flux balance analysis (FBA) by using MitoCore’s default parameters that have been experimentally recorded for heart tissue. FBA has been described extensively elsewhere, but can summarised as calculating the reaction turnover, or fluxes (flows) of metabolites through a network of biochemical reactions assuming a pseudo steady state [3]. The fluxes through the network are constrained by the stoichiometry and directionality of the reactions as well as flux capacity and cytosol boundary uptake ranges. Cytosol boundary transport steps model the import and export of metabolites to the cell, but the overall rate of production and consumption of metabolites is assumed to be zero, hence a pseudo steady state. A metabolic objective function is chosen for a simulation, and FBA is used to calculate an optimal set of reaction fluxes that maximises this function. For this simulation we use maximum ATP production as the objective function because energy generation is one of the primary purposes of central metabolism in cardiomyocytes.

For simulations of ATP yield, all cytosol boundary uptake fluxes for metabolites that could be degraded to produce ATP were set to zero, oxygen was increased to 50 μmol/minute/gram of dry weight (so that limited oxygen availability would not affect the results), while other cytosol boundary conditions were unaltered. The uptake flux of each metabolite of interest was then increased to 1 μmol/minute/gram of dry weight.

For simulations of proton leak, the lower bound of the reaction representing the gene UCP2 (Reaction ID: HtmB_MitoCore) was increased over a series of simulations, thus forcing a minimum flux through the reaction.

All FBA simulations were performed using MATLAB (Math Works, Inc, Natick, MA) and the COBRA Toolbox [38], with the linear programming solver GLPK (http://www.gnu.org/software/glpk).

## Availability of data and material

All data generated or analysed during this study are included in this published article and its supplementary information files. In addition, the model and annotation file will be available at the MRC Mitochondrial Biology Unit website (http://www.mrc-mbu.cam.ac.uk/mitocore/).

## Author’s contributions

ACS and AJR conceived the study. ACS and JPM devised the methodology. ACS created the model.

ACS, FE, JPM, and AJR participated in the curation and testing of the model.

All authors contributed to, read and approved the final manuscript.

## Acknowledgements

We wish to thank Lukasz Zielinski and Alexander Smith for their input and testing of the model, and Edmund Kunji for discussions on the transport of metabolites across the mitochondrial membrane.

## Supporting information captions

Additional file 1. MitoCore model encoded in the SBML format. (XML 795KB)

Additional file 2. Companion annotation spreadsheet recording evidence and provenance for reactions and parameters used in MitoCore and fluxes using default parameters for cardiomyocytes. (XLSX 247 KB)

Additional file 3. Flux distributions around the MitoCore model for maximum ATP production with different metabolic fuels. (XLSX 263 KB)

Additional file 4. Flux distributions around the MitoCore model for maximum ATP production with varying proton leak through reaction representing UCP2. (XLSX 230 KB)

Additional file 5. Experimental measurements of the relative contributions of ΔpH and ΔΨ to the proton motive force. (XLSX 12 KB)

Additional file 6. Table of 105 new or altered reactions from those in the Virtual Metabolic Human database (Recon 2). (XLSX 85 KB)

